# Testing hypotheses of skull function with comparative finite element analysis: three methods reveal contrasting results

**DOI:** 10.1101/2024.11.20.624579

**Authors:** D. Rex Mitchell, Stephen Wroe, Meg Martin, Vera Weisbecker

## Abstract

Comparative finite element analysis often involves standardising aspects of the models to test equivalent loading scenarios across species. However, in the context of feeding biomechanics of the vertebrate skull, what is considered “equivalent” can depend on the hypothesis. We use 13 skulls from diverse group of marsupial bettongs and potoroos (Potoroidae) to demonstrate that that scaling muscle forces to standardise unique aspects of biting mechanics can produce contrasting results of comparative stress or strain that are differentially suited to test specific kinds of hypotheses. We propose three categories of hypotheses for skull biting mechanics which each involve a unique method of muscle scaling: those comparing (1) the skull’s efficiency in distributing input muscle forces, via standardising input muscle force to size, (2) morphological biting adaptation through standardising mechanical advantage to simulate size-independent, equivalent bites, and (3) feeding ecology affected by size, such as niche partitioning, via standardising bite reaction force.

**SUMMARY STATEMENT:** Common approaches for scaling muscle forces in skull finite element models might not always offer reliable results for all hypotheses. We provide a framework for selecting the appropriate method.

## INTRODUCTION

Comparisons of mechanical performance provide valuable insights into how anatomical structures have evolved in response to functional demands. These not only reveal how species have specialised to fulfill their ecological roles, but also shed light on evolutionary relationships and biomechanical trade-offs (Pérez-Ramos et al., 2020; Sansalone et al., 2024). As often the first point of contact between an animal and its sustenance, mechanical performance of the feeding system in particular is an important determinant of an animal’s relative fitness and survival (Ross and Iriarte-Diaz, 2014).

Over the last few decades, computer-assisted biomechanical methods and their application to the study of skeletal function have provided researchers with new tools to test hypotheses related to dietary adaptation. In particular, finite element analysis (FEA), a computational engineering technique used to simulate the behaviour of geometrically complex objects under load, has become established as a powerful tool in modelling the relationship between shape and function in the vertebrate skeleton (e.g., Bright, 2014; Korioth and Versluis, 1997; Rayfield, 2007; Wroe, 2010). FEA simplifies complex structures that are difficult, or otherwise computationally impossible, to assess, through discretisation into a large number of simple objects (elements). These elements, joined at their corners (nodes) to form a contiguous mesh, can be assigned a range of material properties. Simulations of loading involve the application of external forces and the constraint of specific nodes. With respect to the skull and assessing its ability to bite, simulations of jaw adductor muscle force are applied, while the temporomandibular joints and bite point(s) are constrained. These can then yield joint reaction forces and bite reaction forces from solved models. Other metrics returned by the models include the mechanical advantage, or leverage, of bite force production (input force applied/bite reaction force), strain (i.e., deformation, or bending of the structure, defined as change in length/initial length), and stress (force per unit of area). These are key variables in determining mechanical performance and useful for quantifying morphological adaptation because it is non-invasive and repeatable (Wroe, 2010).

To ensure that the stress and strain values obtained from finite element models (FEMs) accurately represent real-world scenarios, muscle dissections and *in vivo* validations of strain magnitudes are obtained from organic samples for comparison. Such validations are difficult to obtain for many species, and impossible for extinct species. To allow for this uncertainty, a simplified comparative approach can be used for non-clinical research whereby the material properties applied to elements are often homogeneous and the simulated forces are arbitrarily defined but then standardised for all models to have consistent ratios of force to size (Dumont et al., 2011a; Dumont et al., 2009). This procedure is not intended to produce mechanical data that reflects biological reality, but instead aims to provide a relative comparison of different species performing the same simulated action. In the context of comparing feeding biomechanics through overall performance of the skull, many studies have shown that allocating homogeneous, isotropic material properties results in negligible differences in broad patterns of stress and strain (e.g., Fitton et al., 2015; Panagiotopoulou et al., 2011; Strait et al., 2010; Walmsley et al., 2013). Through these simplified constructions, lower stress or strain magnitudes in one skull compared to another skull would typically be interpreted as representative of better performance through greater resistance to bending of the bone and potential injury (e.g., Dumont et al., 2011a; Ledogar et al., 2022; Mitchell et al., 2018; Wroe, 2010). Comparative FEA has been used in this way to great effect across diverse extant and extinct vertebrates, particularly to assess feeding biomechanics of the vertebrate skull (Attard et al., 2011; Cook et al., 2021; Cox et al., 2011; Cox et al., 2015; Dumont et al., 2011a; Dumont et al., 2011b; Ferrara et al., 2011; Figueirido et al., 2014; Ledogar et al., 2016; Ledogar et al., 2018; Ledogar et al., 2022; Mitchell, 2019; Mitchell et al., 2018; Mitchell and Wroe, 2019; Oldfield et al., 2011; Rayfield, 2011; Slater et al., 2009; Smith et al., 2015b; Strait et al., 2009; Tseng, 2008; Tseng et al., 2011; van Heteren et al., 2021; Wroe, 2007; Wroe et al., 2013; Wroe et al., 2007). Through these studies, we have gained valuable insights into the relationship between skull morphology and mechanical performance, with applications towards conservation, ecology, evolution, and palaeontology.

The goal of the comparative approach is to standardise parameters as much as possible to subject each model to equivalent, or near equivalent, loading conditions. However, what constitutes equivalency might differ depending on the hypothesis and various standardising regimes used by researchers might not be adequate for answering all possible questions focussed on how well different species perform during biting. For example, research using standardised muscle force on larger samples of models has identified inconsistencies resulting from interspecific variation in mechanical advantage that can impact strain magnitudes (Mitchell et al., 2018). Furthermore, the roles that size and shape both play in overall function of the skull (Chamoli and Wroe, 2011; Mitchell et al., 2024c), suggest that common approaches used to remove variation in size are potentially not appropriate for all questions relating to animal ecology and feeding behaviour.

Because most hypotheses of cranial biomechanics focus on the relationship between shape and performance (i.e., functional morphology), without consideration of how size impacts on cranial performance, variation due to size is recommended to be removed through standardising the ratios of size and muscle force (Dumont et al., 2011a; Dumont et al., 2009; Strait et al., 2009). There are two common methods that aim to achieve this. The first is to scale all models to the same size (usually determined as having the same surface area) and apply identical muscle forces to all models (Attard et al., 2011; Dumont et al., 2009; Wroe et al., 2010). The second is to scale the muscle forces of an arbitrarily chosen reference specimen to the size of all other specimens (McHenry et al., 2007). This is most often done using skull volume to a two thirds power (Cook et al., 2021; Ledogar et al., 2016; Ledogar et al., 2018; Mitchell, 2019; Smith et al., 2015a; Smith et al., 2015b; Strait et al., 2010). Either approach of standardising skull size and muscle force ratios is intended to provide a means of comparing the ability of skull shapes to convert muscle force to bite force. However, standardising muscle force is just the first of three standardisations that can be enacted on the jaw lever system in FEA. The other two are standardising mechanical advantage and standardising bite force. Standardising each of these might produce different comparisons of stress or strain, especially if the sample has appreciable variation in mechanical advantage or size.

In studies of feeding biomechanics, mechanical advantage refers to how much muscle force can be converted to bite force. In simplest terms, this is defined by the relative lengths of the in-lever (distance of the average muscle force vector from the temporomandibular joint) and out-lever (distance of the biting tooth from the temporomandibular joint). All else being equal, skulls with orthognathic proportions, or “shorter faces”, therefore tend to transfer more input muscle force to the biting teeth, representing greater mechanical advantage (Goswami et al., 2010; Mitchell et al., 2024c). Selection for mechanical advantage is considered one of many important determinants of mammalian skull diversity (Dumont et al., 2009; Dumont et al., 2014; Ledogar et al., 2022; Mitchell et al., 2024c), with rostrum length playing a key role in how efficient a jaw lever system is. Standardising input muscle forces is certainly reasonable for calculating mechanical advantage and assessing stress or strain of the skull under equivalent muscle loading, but this method might not account for variation in mechanical advantage. Consequently, models of species with lower mechanical advantage might produce weaker bites from the same input muscle force and therefore experience lower stress and deformation throughout the facial skeleton. Because lower strain magnitudes are often interpreted to represent morphology better suited to resist deformation and injury during biting, species with longer faces could risk being interpreted as better adapted to forceful biting than they are. Conversely, skulls with high mechanical advantage would produce stronger bites and therefore experience greater deformation of the facial skeleton than should be expressed during equivalent biting behaviours (Mitchell et al., 2018). Therefore, comparative biomechanics that asks how efficiently different skull shapes produce equivalent bites benefit from standardising of mechanical advantage.

There are two main methods that aim to standardise mechanical advantage. The first is done on models that have been scaled to equivalent size, by rescaling the initial muscle forces to result in a common bite force (Fig. 1A) (Attard et al., 2011; Oldfield et al., 2011; Wroe et al., 2010). The goal is to ensure that all models are performing an equivalent action, for their size, and the stress and strain magnitudes therefore adequately reflect an equivalent action of the jaw, when adjusted for size. A second method can be done on models of their natural size that have had their initial muscle forces scaled to volume to the two thirds power. This involves rescaling initial input muscle forces to produce bite reaction forces expected from a standardised mechanical advantage (Fig. 1B) (Mitchell et al., 2018; Mitchell and Wroe, 2019). For the many taxa that exhibit an increase in face length with increased size (a CREA pattern) (Cardini et al., 2015; Cardini and Polly, 2013; Mitchell et al., 2024b; Mitchell et al., 2024c; Tamagnini et al., 2017), mechanical advantage will often decrease with increasing size. The predictive regression line of standardised mechanical advantage would therefore follow a path of increasing bite reaction force when compared to the initial loadings, indicating that larger individuals would need greater muscle forces applied to achieve equivalent relative biting actions (Fig. 1C). Both of these methods for standardising mechanical advantage aim to account for variation in mechanical advantage and might therefore more accurately compare the performance of the skull for hypotheses relating to skull shape and biting adaptation. However, these approaches are not necessarily adequate for hypotheses of a more ecological focus, such as how well animals of different sizes can bite the same piece of food. Here, we propose that such hypotheses also might benefit from a third standardising technique that includes size variation in the analysis. This involves the standardisation of absolute bite force.

**Figure 1:**
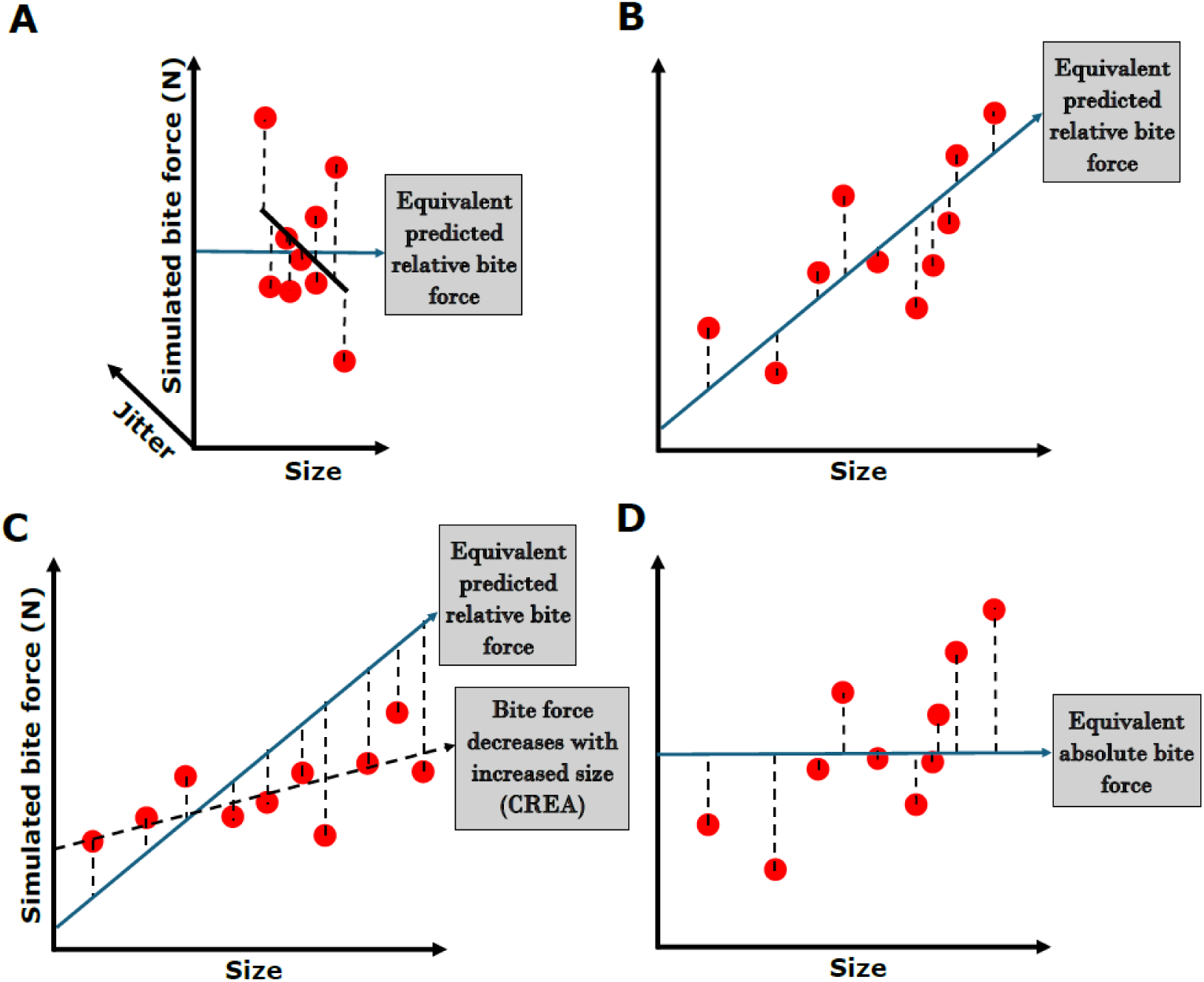
Methods of muscle force scaling in comparative FEA. (A) When models initially scaled to the same size have the same input muscle forces applied, all bite reaction forces plot on a single point of the size axis but show variation due to mechanical advantage (jitter variation on a z-axis added for clarity). Muscle forces can be rescaled such that each model produces the same bite reaction force. Since models have been scaled to the same size, this approach aims to simulate relative (size-independent) bite force. (B) When models retain their natural size and instead have muscle forces scaled to size, bite reaction forces from initial simulations are plotted against size. The mean mechanical advantage of the sample is multiplied with initial muscle forces to predict equivalent relative bite force. Initial muscle forces can then be rescaled to produce the same relative bite force. (C) When samples follow a CREA pattern of decreasing mechanical advantage with size, standardising mechanical advantage requires greater input muscle force applied to larger species and reduced input muscle force applied to smaller species. (D) To factor size into function for hypotheses relating to ecology, muscle forces on models with original size retained can also be rescaled to produce the same absolute bite force. In this case, we use the mean bite force from the sample, but the chosen bite force is arbitrary.

Dumont et al. (2009) advised the disentangling size and shape for comparative finite element modelling. Their perspective came from a foundational prioritisation of shape as the predominant indicator of function and emphasised the importance of shape differences in relation to mechanical performance. However, size is a crucial determinant of an animal’s skull structure and absolute function in an ecosystem (Alexander, 1985; Chamoli and Wroe, 2011; Doube et al., 2011; Mitchell et al., 2024c). Adaptive trade-offs in the mammalian cranium across size ranges might suggest that size and morphology should not be separated when considering adaptation to environmental parameters that can oftentimes remain constant regardless of size (Marcy et al., 2024; Mitchell et al., 2024b; Mitchell et al., 2024c). In particular, all else being equal, if larger species can bite harder than smaller species owing to their larger skulls, jaw muscles, and teeth, the skull of a larger species will more easily bite into an object of a given size and composition than the skull of a smaller species. This potentially allows the evolution of lower mechanical advantage in larger species (e.g. through elongation of the rostrum) without sacrificing absolute bite force (Mitchell et al., 2024c).

Because both size and shape jointly determine the mechanical capabilities of an animal under natural settings, we suggest that size should be considered in hypotheses relating to comparisons of behavioural ecology, competition, niche differentiation, and overall potential interactions and impacts on the environment. This would involve scaling muscle forces, on models of their natural size, to produce the same absolute bite force (Fig. 1D). Surprisingly, this approach has almost never been taken (but see Fitton et al., 2015). But by factoring size into the scaling protocol of FEA, thus representing absolute (size-dependent) bite force comparisons, we would expect to see strain magnitudes across models that can compare quite differently to those offered by relative (size-independent) bite force simulations. For example, smaller skull models should exhibit greater deformation, while larger models should exhibit reduced deformation compared to size-independent simulations on the same models.

Here, we show that all three standardising approaches produce different estimations of cranial stress or strain (stress and strain are directly proportional when using homogeneous, isotropic models). We chose the family of Potoroidae, including Bettongs and potoroos, as an ideal group to compare results from the three methods. Potoroids exhibit a suitably large range of elongation of the facial skeleton that appears to be independent of size, suggesting strong selection on mechanical advantage (Mitchell et al., 2018). All extant species specialise on highly digestible foods with discrete and constant mechanical properties, such as fungal sporocarps (truffles), roots, tubers, seeds, nuts, fruits, and insects, and comprise the diets of each species to varying proportions (Mitchell et al., 2024a; Seebeck et al., 1989). Importantly, species with greater focus on softer fungi tend to have longer snouts, while shorter faces tend to be accompanied by diets with harder foods, such as nuts, seeds, and roots (Mitchell et al., 2024a; Mitchell et al., 2018). To demonstrate the predictive power of our approach, we also include an extinct species*, Caloprymnus campestris*, which had a small skull for a potoroid, but particularly robust craniofacial proportions. Specifically, we aim to test three hypotheses:

1. when applying only the initial standardised input muscle forces, variation in mechanical advantage can confound strain magnitudes for assessments of equivalent biting ability, such that species with long faces will have lower strain and short-faced species will have higher strain. This would be contrary to what should be expected from their diets; (2) when muscle forces are rescaled to standardise mechanical advantage, and therefore produce equivalent bite forces relative to size, skulls with shorter snouts (i.e., greater advantage) will exhibit lower strain than skulls with longer snouts during equivalent bites; and (3) when rescaling muscle forces to standardise absolute bite forces, the model of the smaller, more robust cranium of *C. campestris* will experience higher mean strain than other species with similarly efficient skulls, suggesting that it was not as capable of biting the same foods as larger species with similarly robust skull anatomy.

## MATERIALS AND METHODS

Our sample included thirteen intact, adult skull specimens from the South Australian Museum and the Queensland Museum: 2 x *Aepyprymnus rufescens* (SAM M2751 & SAM M12026), 2 x *Bettongia gaimardi* (SAM M7384 & SAM M7389), 2 x *Bettongia lesueur* (SAM M1705 & SAM M18492), 2 x *Bettongia penicillata* (SAM M8285 & SAM M11252), 2 x *Bettongia tropica* (QM JM10030 & QM JM12495), 2 x *Potorous tridactylus* (SAM M7381 & SAM M9013), and 1 x *Caloprymnus campestris* (SAM M3257). We micro-computer-tomography (μCT) scanned all specimens using the Flinders Tonsley Nikon XT H 225ST CT Scanner (Large-Volume Micro-CT System), with isotropic pixel sizes ranging from 20-25 µm.

We generated 3D surface meshes of the crania and mandibles from the μCT data in Mimics (Materialise v.26.0). For each model, the cranium was centred and then oriented to align the temporomandibular joints and dental arcades with the Y-axis for restraining. The mandible was then positioned for near occlusion at the incisors to best simulate biting a smaller-sized object.

We exported the cranial meshes and converted to them to FEMs (volume meshes) using 3-Matic (Materialise v. 18.0). Each model consisted of approximately 1-2 million 3D tetrahedral elements (bricks). Models were then imported into Strand7 (v. R3.1.3.a) for finite element modelling. We assigned homogeneous, isotropic material properties of average mammalian bone (Young’s modulus: E ∼ 20 GPa; Poisson’s ratio: v = 0.3) (Figueirido et al., 2014; Mitchell, 2019; Mitchell et al., 2018; Mitchell and Wroe, 2019; Sharp, 2015; Tseng et al., 2011). Homogeneous and isotropic material properties were considered acceptable to assess the relationship between gross cranial morphology and biting performance (Fitton et al., 2015; Strait et al., 2010; Walmsley et al., 2013). As is common practice in comparative FEA studies (Dumont et al., 2005; Mitchell, 2019; Wroe et al., 2007), we emphasise that our results should be considered in a relative context and not as actual *in vivo* strain magnitudes.

We partitioned major jaw adductor muscle groups (masseter, temporalis, and pterygoids) using overall proportions for *Potorous (Warburton, 2009)*. We sub-divided the proportions of the masseter complex (superficial, intermediate, and deep) and pterygoids (medial and lateral) using subunit proportions of a red-necked wallaby (Mitchell et al., 2018). A total muscle force of 100N was partitioned according to these muscle proportions and a specimen (*A. rufescens* M2750) was randomly chosen (alphanumerically) as the reference specimen to scale muscle forces for all other models. The absolute reference muscle force is arbitrary and does not change the predicted distribution of stress or strain in a structure (Mitchell, 2019; Wroe, 2007; Wroe et al., 2007). The muscle forces were scaled to cranial volume using a 2/3 power rule (Strait et al., 2010) (Table S1).

Muscle plates representing origins and insertions were defined using Geomagic (v.2021). The masticatory muscle forces were applied to cranial plates using BoneLoad (Davis et al., 2010; Grosse et al., 2007). This software orients the forces from the cranial muscle origins to the centroids of their respective insertions, following the curvature of the bone. The loaded plates were imported into Strand7 and zipped to the nodes of their corresponding elements. A single node on the tip of each biting tooth was restrained against translation in the vertical axis: both incisors for a bilateral incisor bite, and a single, right side P3 premolar or M2 molar for unilateral simulations. A single node at each temporomandibular joint was restrained against translation for all axes. The models were then solved as linear static.

From these initial simulations, we extracted reaction forces from the biting teeth. We then calculated mechanical advantage by dividing the bite reaction force by the input muscle force. To address hypotheses relating to morphological biting adaptation, we rescaled the muscle forces to produce equivalent bite forces, relative to size (Tables S2-S4). To do this, we first calculated the expected bite force for each specimen by multiplying the input muscle forces by an identical value of mechanical advantage, arbitrarily represented by the mean mechanical advantage of all models (Tables S2-S4). Dividing this predicted bite force by the initial bite reaction force of the simulation gives the scaling factor to multiply all initial muscle forces by. The scaling factor is therefore given by the following equation:

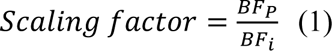

Where *BF_p_* is the predicted bite force scaled from the mean mechanical advantage of all models and *BF_i_* is the bite reaction force of the initial simulation. We then multiplied all initial muscle forces by the scaling factor. This method is similar in aim to rescaling the models to the same size and applying muscle forces that result in the same bite reaction force for a given size (e.g., Attard et al., 2011; Oldfield et al., 2011; Wroe et al., 2010); however, we chose this approach because it allows for the same models to be used in all our scaling scenarios. This procedure corrected for the effects mechanical advantage on strain magnitudes during equivalent bites. Differences in mean strain under these conditions should therefore only reflect differences in how well a given cranial morphology deforms, or bends, during a given bite reaction force, relative to size (Dumont et al., 2009; Mitchell et al., 2018; Smith et al., 2015a).

For hypotheses concerning the comparative ability of different animals to withstand identical bites in the wild, we needed the models to produce equivalent absolute bite forces (Tables S5-S7). In this case, predicted bite forces were standardised to the mean value of all bite reaction forces from the initial simulations (Fig 1D). The precise value used for equivalent absolute bite force is arbitrary in a comparative context, but using the mean bite reaction force allowed us to use the same colouration scales in all visualisations. All muscle forces and rescaled muscle forces applied to all models are available in the supplementary information (Tables S2-S7). The scaling factor for this method is given by the same equation as above, but with the predicted bite force being the mean bite reaction force from the initial simulations. We emphasise that the only changes made to the models across these scaling approaches is the amount of muscle force applied.

We used von Mises (VM) strain to represent deformation of all models and visualised this using VM heatmaps, set to a threshold of 15µε. However, the comparative results obtained can also hold true for stress in homogeneous, isotropic models, because stress and strain are directly proportional under such conditions. We extracted strain magnitudes from all elements of each model and created custom R code (R Development Core Team, 2023) to automatically remove the top 1% of strain values, because these typically relate to artifactual inflated regions close to the restraints (Marce-Nogue et al., 2017; Walmsley et al., 2013). We plotted histograms of each simulation that depicted mean VM strain for each model. A higher degree of mean strain indicates greater deformation of the model during the simulated activity.

## RESULTS

Potoroos and bettongs had a high degree of variation in mechanical advantage (Fig. 2). Across all simulations, the highest mechanical advantage was seen in *B. lesueur*, closely followed by *C. campestris*. The lowest mechanical advantage was seen in *P. tridactylus*. This variation in mechanical advantage means that under the initial muscle loading scenarios, all species were producing bites of different magnitude, relative to their size.

**Figure 2:**
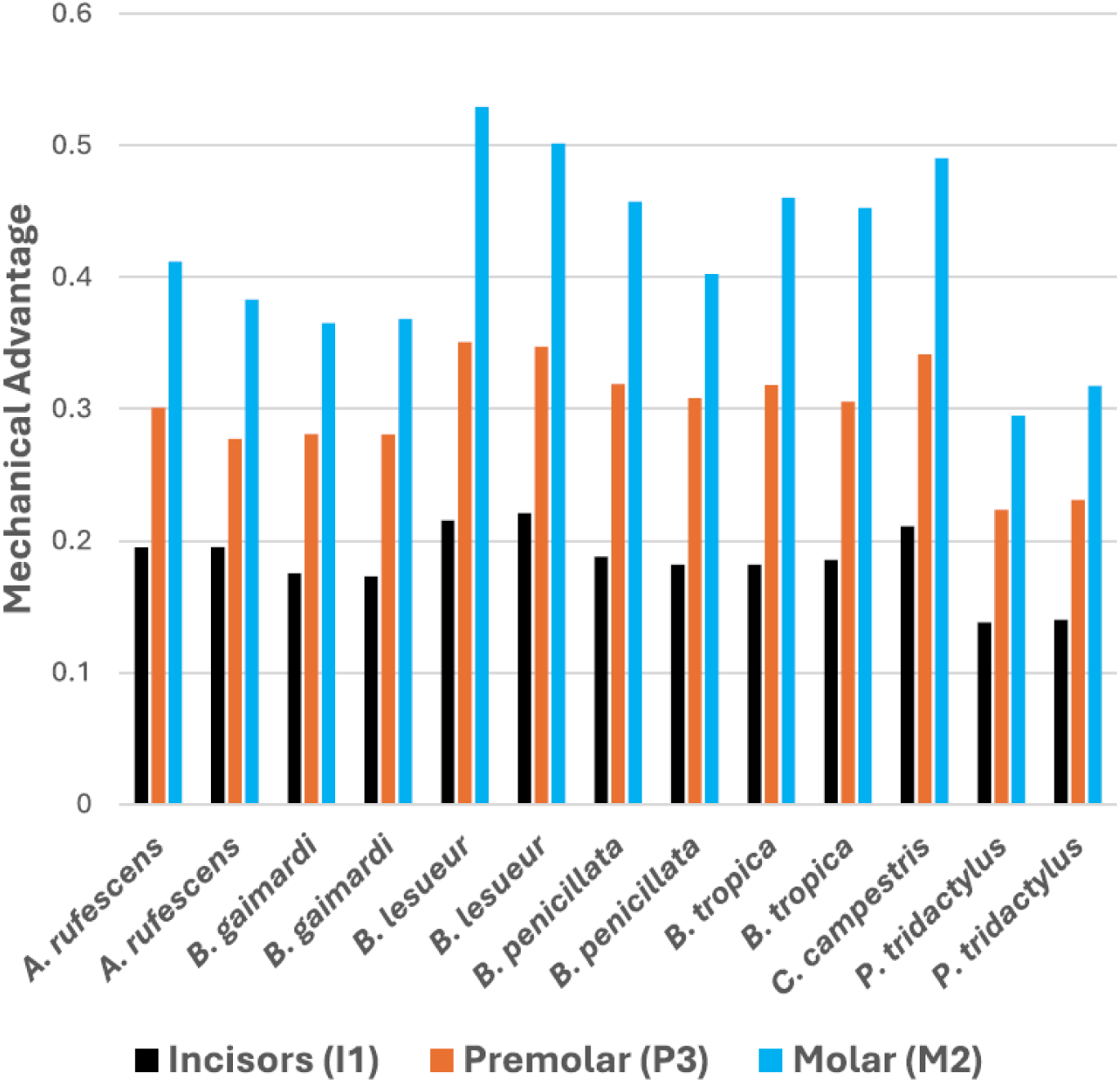
Mechanical advantage across all simulated bite scenarios.

Dorsal and lateral views of VM strain for all 117 simulations are presented in Supplementary Figure S1. However, we present some examples here to highlight contrasting results in comparative strain magnitudes. As predicted, lower mechanical advantage in *P. tridactylus* resulted in lower deformation under initial loading conditions with standardised muscle force. Rescaling the muscle forces to produce equivalent relative bite forces (standardised mechanical advantage) both decreased the strain in models with high advantage and increased the strain of models with low advantage. This was particularly obvious *in P. tridactylus* (Fig. 3 and Fig. 4), which needed up to 42% more input muscle force to match the predicted relative bite forces of the other species (Tables S1-S7).

**Figure 3:**
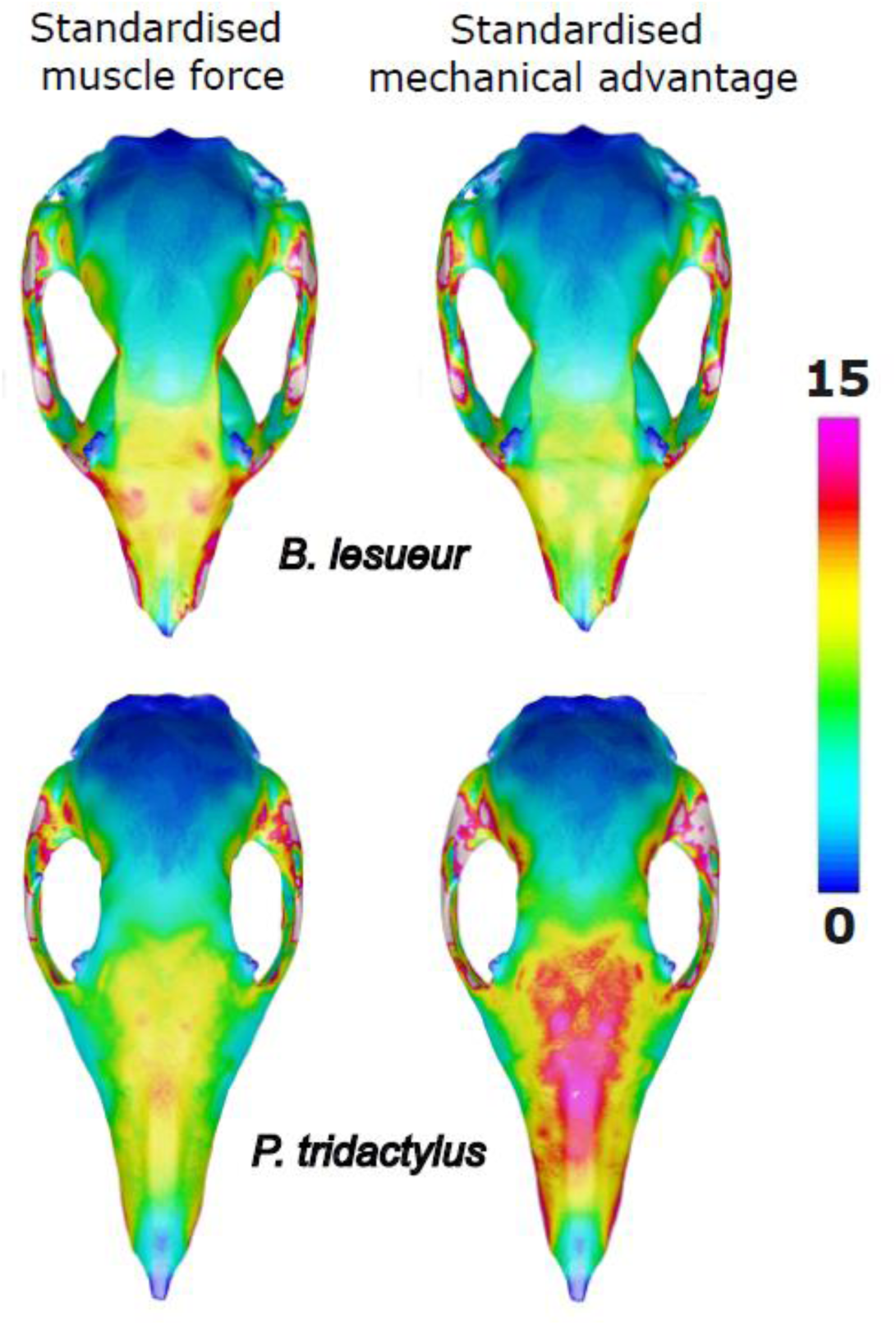
Simulations with muscles (A) representing equivalent relative input muscle force through scaling to cranial size, (B) rescaled to produce equivalent relative bite force through standardised mechanical advantage.

**Figure 4:**
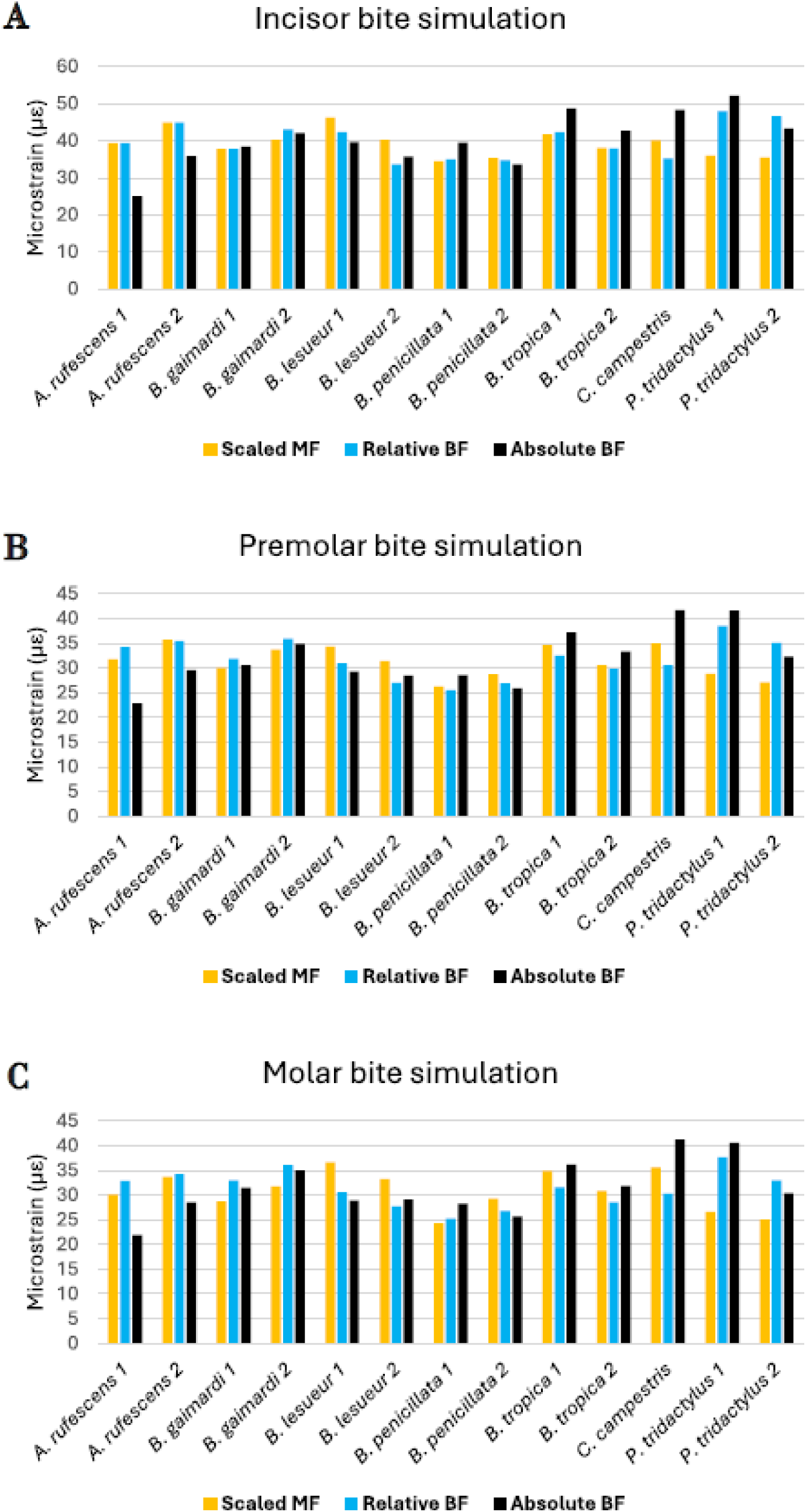
Mean element strain of all simulations: (A) incisor bite, (B) P3 premolar bite, (C) M2 molar bite. MF = muscle force, BF = bite force.

These changes in applied forces reversed the comparative patterns of mean strains (Fig. 4), such that the most deformation (indicated by mean brick strain) was then observed in *P. tridactylus*, while the least was seen in *C. campestris*, *B. penicillata*, and one of the *B. lesueur*. The overall patterns across incisor, premolar and molar bite simulations were mostly consistent, however, *C. campestris* performed better during incisor biting. Among *Bettongia*, higher mean strains were consistently found in *B. gaimardi*, *B. tropica*, and one of the *B lesueur*.

Rescaling the muscle forces to produce equivalent absolute bite forces gave similar results for most species, with the high mean strain still found in *P. tridactylus*. But there was a noticeable shift in the performance of much smaller *C. campestris*, which this time experienced similar mean strain to *P. tridactylus* under this loading regime (Figs 4 and 5). Fig. 4 also identifies noticeable intraspecific variation in mean strain relating to size variation of the specimens, particularly in *A. rufescens* and *P. tridactylus*; however, these do not appreciably impact overall interspecific comparative patterns.

**Figure 5:**
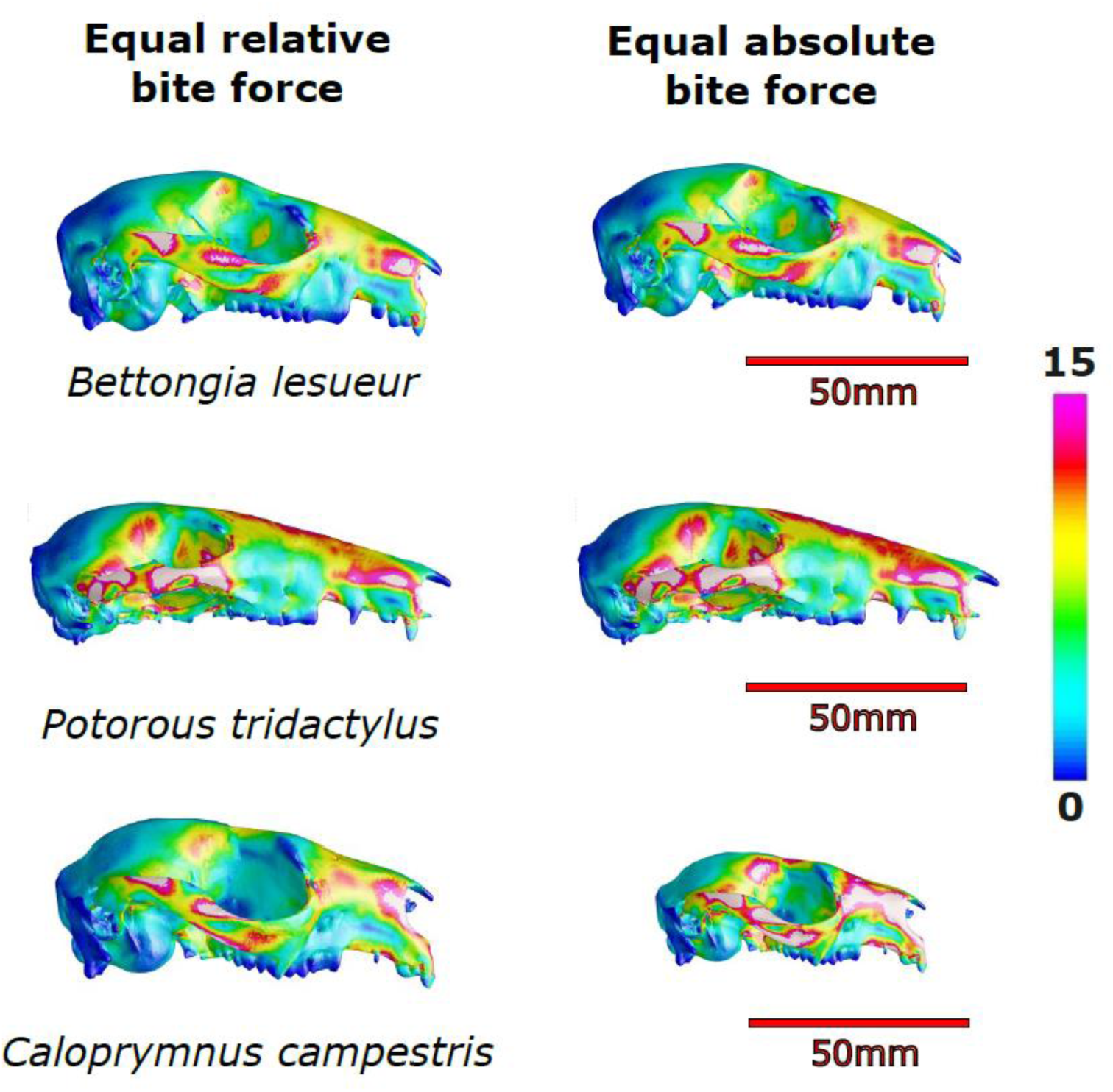
Simulations with muscles (A) scaled to produce equivalent relative (size-independent) bite force, (B) scaled to produce equivalent absolute (size-dependent) bite force.

## DISCUSSION

In this study, we focussed on methods of jaw muscle scaling in comparative finite element analysis of skulls. We highlighted three standardisations in biting comparisons that each require unique approaches to muscle scaling: input muscle force, mechanical advantage, and bite force. Using a sample of potoroid marsupial skulls with appreciable variation in mechanical advantage and size, we show here that each method produces unique results of comparative mean strain, but all are informative for specific cases. Based on our results, we propose that studies aimed at answering questions in the area of feeding biomechanics can be partitioned into three major categories, which each require a specific method of muscle scaling: those interested in (1) bite force production (system efficiency), through standardising input muscle force to cranial size; (2) relative bite force (biting adaptation), via standardising mechanical advantage on models scaled for size-independence; or (3) absolute bite force (feeding ecology) by standardising bite reaction force. Standardising one of these three conditions (muscle force, mechanical advantage, or bite force) produces variation in the other two conditions, potentially leading to differential results of comparative stress or strain. For many taxa studied using comparative FEA, as is the case for several species in this study, the differences between the methods used might be negligible. However, if there are appreciable differences in mechanical advantage or size between the species being modelled, we show that results for the three methods can deviate from each other. We therefore suggest that each of these three major categories might be best examined with their own scaling method.

Importantly, methods 2 and 3 require an initial set of simulations for calculating mechanical advantage and rescaling of muscle forces. Standardising of mechanical efficiency (category 2 method) requires the initial simulations to first be scaled to cranial size (category 1 method). This is to maintain size-independent muscle scaling across specimens, upon which variation in bite reaction force due to variation in mechanical advantage is removed. Without this crucial step, bite reaction forces predicted from standardised mechanical efficiency will not represent size-independent assessments of shape performance. However, the initial input muscle forces used for standardising of bite reaction forces (category 3 method) are arbitrary for subsequent rescaling, because the predicted bite reaction forces are constant and not influenced by size variation.

The initial simulations, with size-standardised input muscle forces applied, provide an estimation of how efficiently different skulls convert a given amount of muscle force to bite force. This method was useful in this study to obtain the mechanical advantage values for rescaling in the two subsequent methods. But our simulations also showed that lower strain magnitudes were found in species with low mechanical advantage. With the understanding that lower strain is an indication of improved bite force capacity (Dumont et al., 2011a; Mitchell, 2019; Wroe, 2010), use of this scaling method might have led to a conclusion that longer-faced species – such as the long-nosed potoroo in our sample - are better adapted for forceful biting.

However, when rescaling muscle forces to produce equivalent relative bite forces, through standardising variation in mechanical advantage, the patterns of the initial simulations were reversed, with the least efficient skulls exhibiting higher mean strain, as would be expected for the longer faced species that feed on high proportions of soft fungi. This indicates that the initial scaling method alone is likely not the most appropriate approach to compare stress or strain magnitudes relating to biting adaptation, owing to the confounding effects of mechanical advantage on bite reaction forces. Hypotheses based on comparing how well skulls disperse a given input force to the joints, teeth, and throughout the overall structure do, however, still exist. For example, more clinically oriented questions relating to stress or deformation experienced by the facial skeleton under specific loading (e.g., for dentistry or facial prosthetics) can be best answered with the initial muscle scaling method. These kinds of questions fall under the (1) bite force production category. However, interspecific studies using comparative FEA of skulls tend to be more interested in how skull shape influences biting performance (category 2). We showed that these questions are better assessed by rescaling muscle forces to produce equivalent relative bite forces through standardising of mechanical advantage.

We compared models producing equivalent relative bite forces by standardising mechanical efficiency to specifically assess the importance of skull shape in biting ability (category 2 hypothesis). Under these conditions, the species with higher mechanical advantage tended to experience lower strain and species with lower mechanical advantage tended to produce greater mean strain magnitudes, when compared to their initial simulations. This reflects the need for skulls with lower advantage to provide more input muscle force to produce equivalent bites, relative to size (Mitchell et al., 2018; Mitchell et al., 2024c). Accordingly, greater stress and strain will be dispersed throughout the facial skeleton. In contrast, skulls with greater mechanical advantage need less muscle force to produce equivalent bites, so mean strain will be lower.

Simulations with standardised input muscle force revealed that the extinct *C. campestris* had a highly efficient skull for transferring muscle force to bite force. Simulations of standardised mechanical advantage further indicated that the species had a skull shape well suited to incisor biting, with strain magnitudes similar to other efficient skulls of *Bettongia* spp. Operating under the assumption that shape alone is indicative of functional adaptation, we might conclude that this species was adapted for more forceful biting and consider the results an indication of a diet high in mechanically resistant foods. However, to compare the ability of the skulls to accommodate a specific bite in an ecological context (category 3 hypothesis), we then rescaled muscle forces to produce equivalent absolute bite forces, simulating each individual biting a similar food item. Under these conditions, *C. campestris* performs considerably worse than other species with similarly efficient skulls, with mean strain more comparable to the mechanically inefficient skulls of *P. tridactylus*. This suggests that the compact appearance and high mechanical advantage of the skull of *C. campestris* are not adaptations for more forceful biting in an absolute sense, but instead compensate for its considerably smaller size. This would allow it to feed on more mechanically similar foods to larger species (Mitchell et al., 2024b), but not to the extent of specialising on particularly resistant foods.

*Caloprymnus campestris* is considered to have been largely plant-eating (phytophagous) (Finlayson, 1932), but remains of beetles have also been found in gut contents (Flannery and Kendall, 1990). Our results thus support a phytophagous diet previously identified for this species, but further predict that *C. campestris* likely had a diet dominated by softer, higher-quality plant material, such as forbs and fresh leaf material. Supplementing the diet with insects is also probable given this is common among potoroids. The higher performance of incisor biting and lower performance of premolar biting in this species suggests greater importance of incisal cropping in this species, rather than premolar biting as is commonly employed for most food processing in other potoroid species (Sanson, 1989). These conclusions would likely have been missed using the other two approaches to scaling muscle forces.

Scaling muscle forces to produce equivalent absolute bite forces also carries some additional considerations. For instance, size influences strain magnitudes within species as well as across species. During bites of equivalent absolute magnitude, the larger specimens of both *A. rufescens* and *P. tridactylus* experience notably lower strain. Intraspecific variation in size should therefore be considered when assessing model performance for simulations of absolute bite force production. This might also open up the methodology to new considerations of how ontogeny, sexual dimorphism, and size variation in general impact niche differentiation and competition.

## ACKNOWLEDGEMENTS

The authors acknowledge the facilities, and the scientific and technical assistance of Microscopy Australia and the Australian National Fabrication Facility (ANFF) under the National Collaborative Research Infrastructure Strategy, at the South Australian Regional Facility, Flinders Microscopy and Microanalysis, Flinders University. We also thank David Stemmer of the South Australian Museum. This research was conducted on the traditional lands of the Kaurna people.

## AUTHOR CONTRIBUTIONS

DRM: conception and design, acquisition of data and analysis, interpretations, and writing; SW: interpretations, and writing; MM: acquisition of data and processing; VW: acquisition of data and analysis, interpretations, and writing. All authors approve of the final version.

## DATA AVAILABILITY

Scans of all specimens are available on Morphosource https://www.morphosource.org/projects/000553610?locale=en

Extracted brick strains and R code are available on Figshare private link: https://figshare.com/s/cf5f814aba28a55c2ad9

