## Supplementary material for "Testing hypotheses of skull function with comparative finite element analysis: three methods reveal contrasting results": Figure S1

**Supplementary Figure 1: Von Mises strain heat maps of all simulations.**

**Incisor bite simulations**

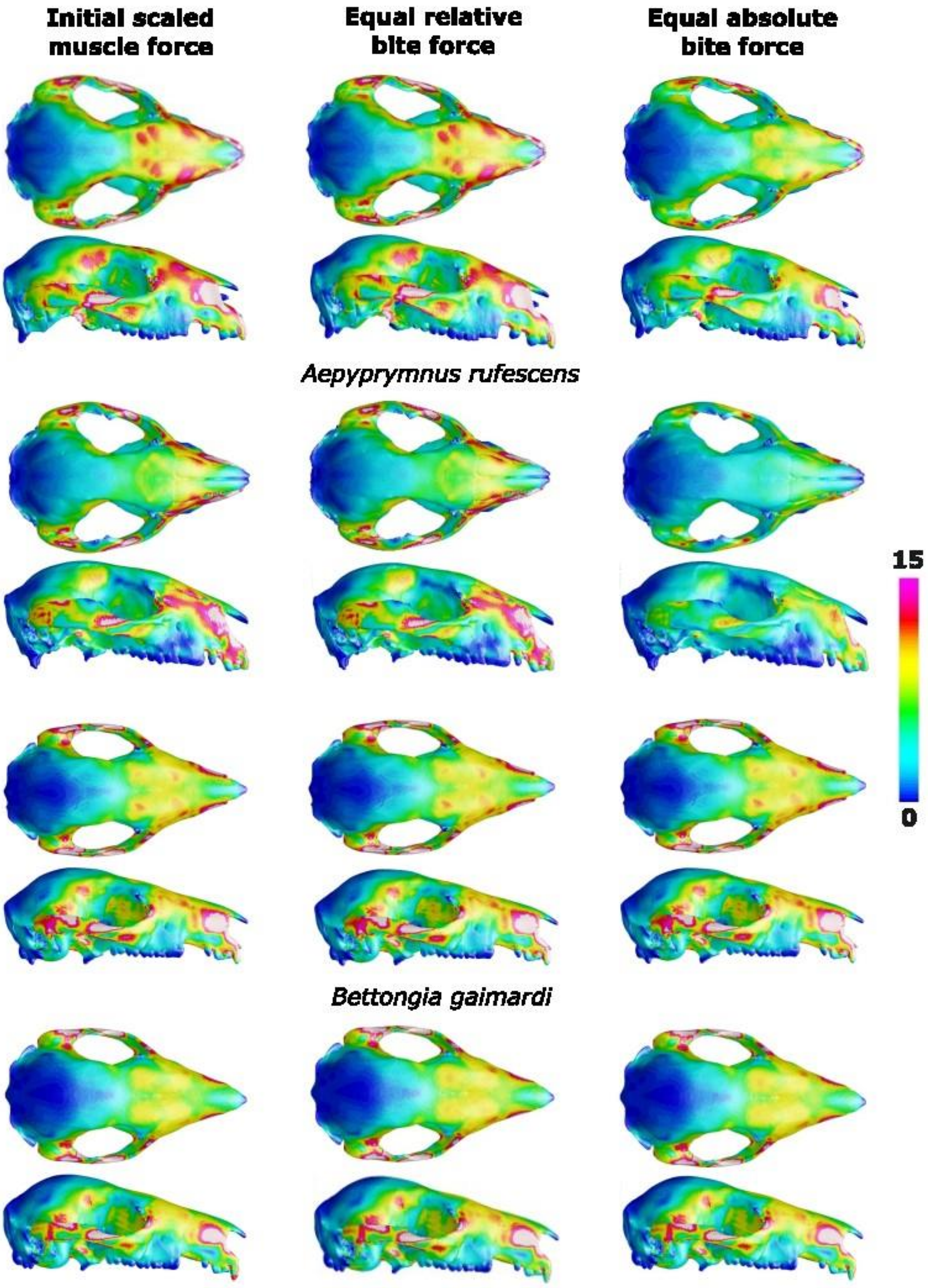

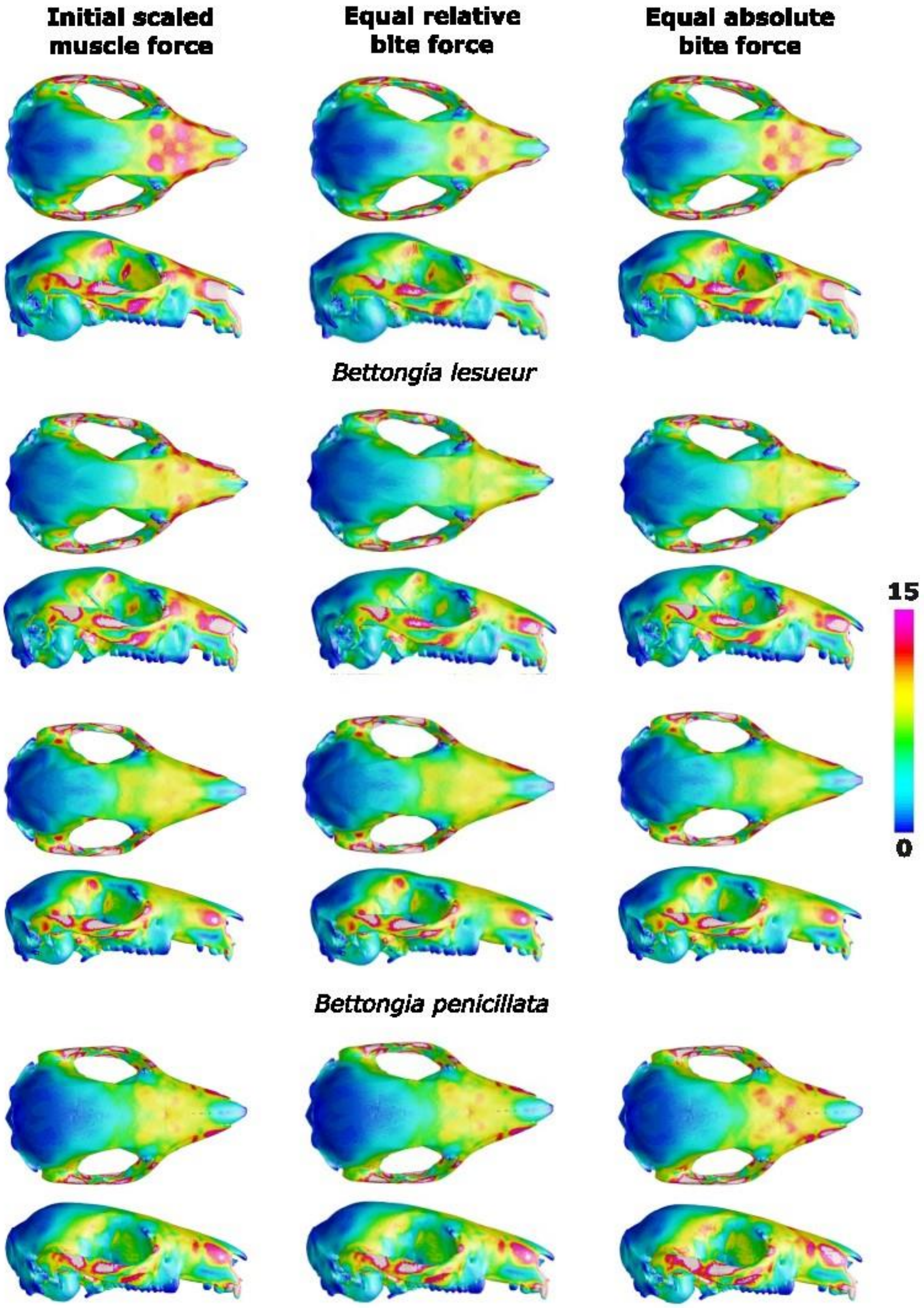
Initial scaled muscle force
Equal relative bite force
Equal absolute bite force
Bettongia lesueur
Bettongia penicillata
15
0

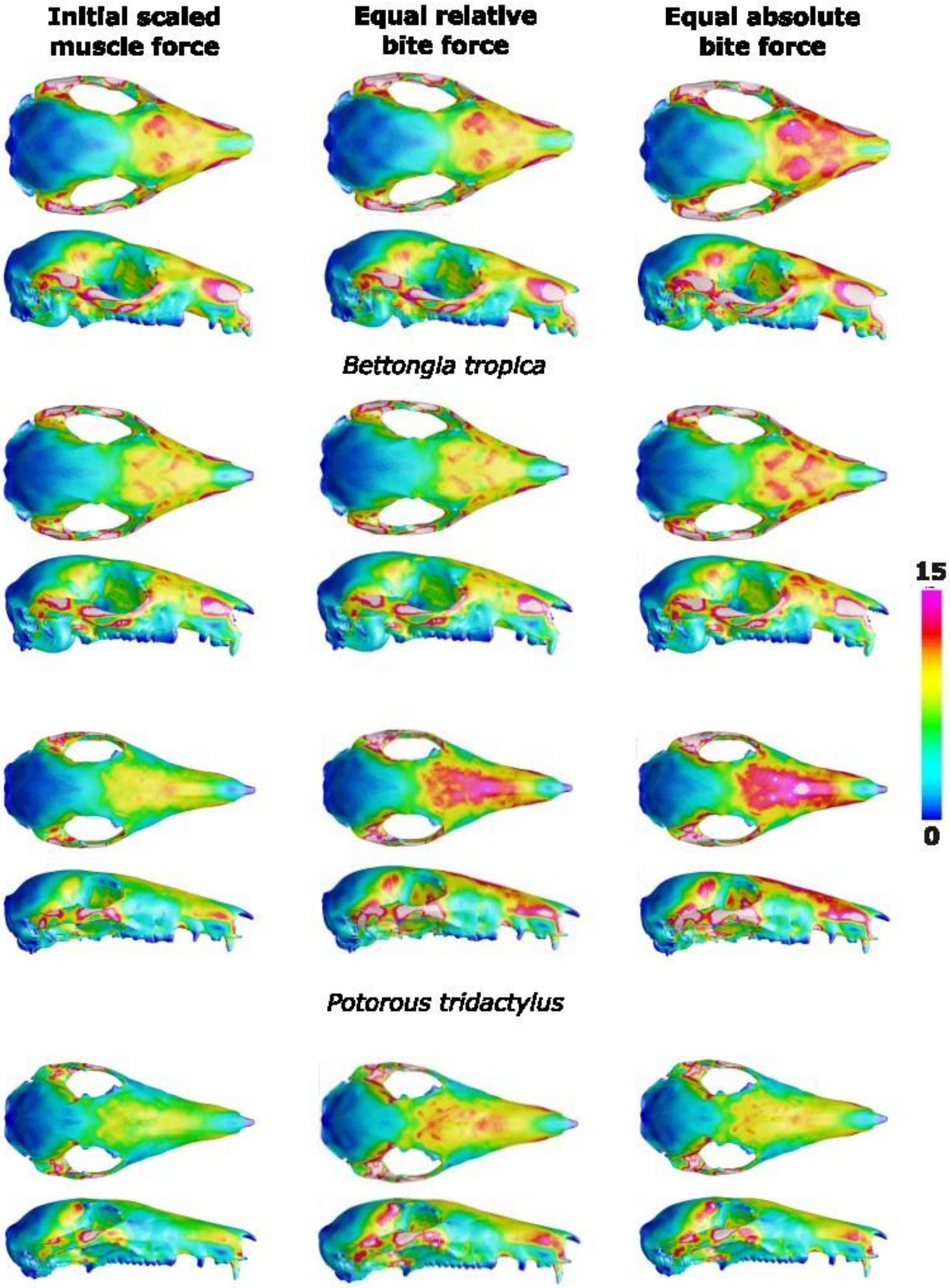

Initial scaled muscle force
Equal relative bite force
Equal absolute bite force
Bettongia tropica
Potorous tridactylus
15
0

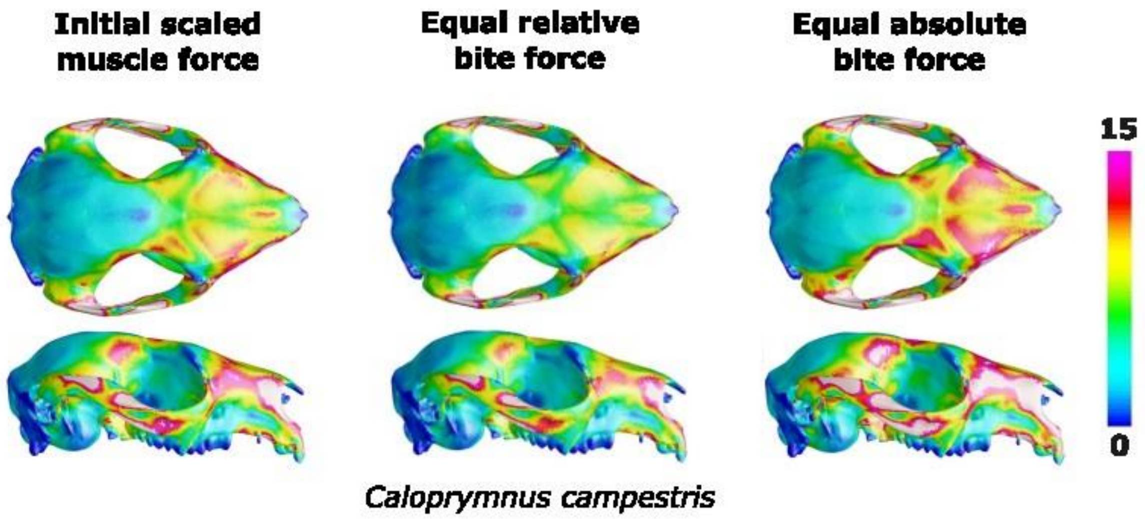
Initial scaled muscle force
Equal relative bite force
Equal absolute bite force
15
0
*Caloprymnus campestris*

**Premolar bite simulations**

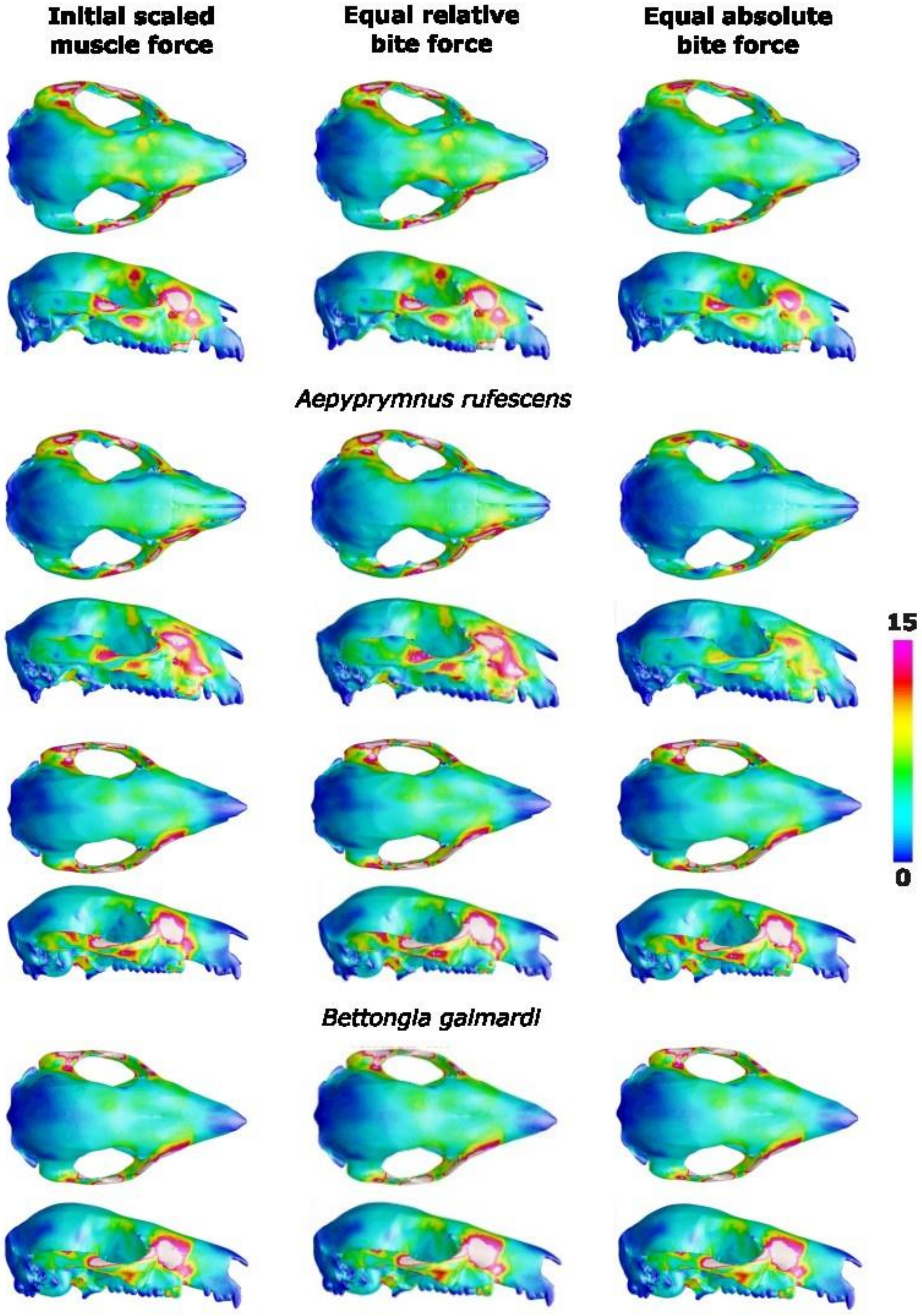

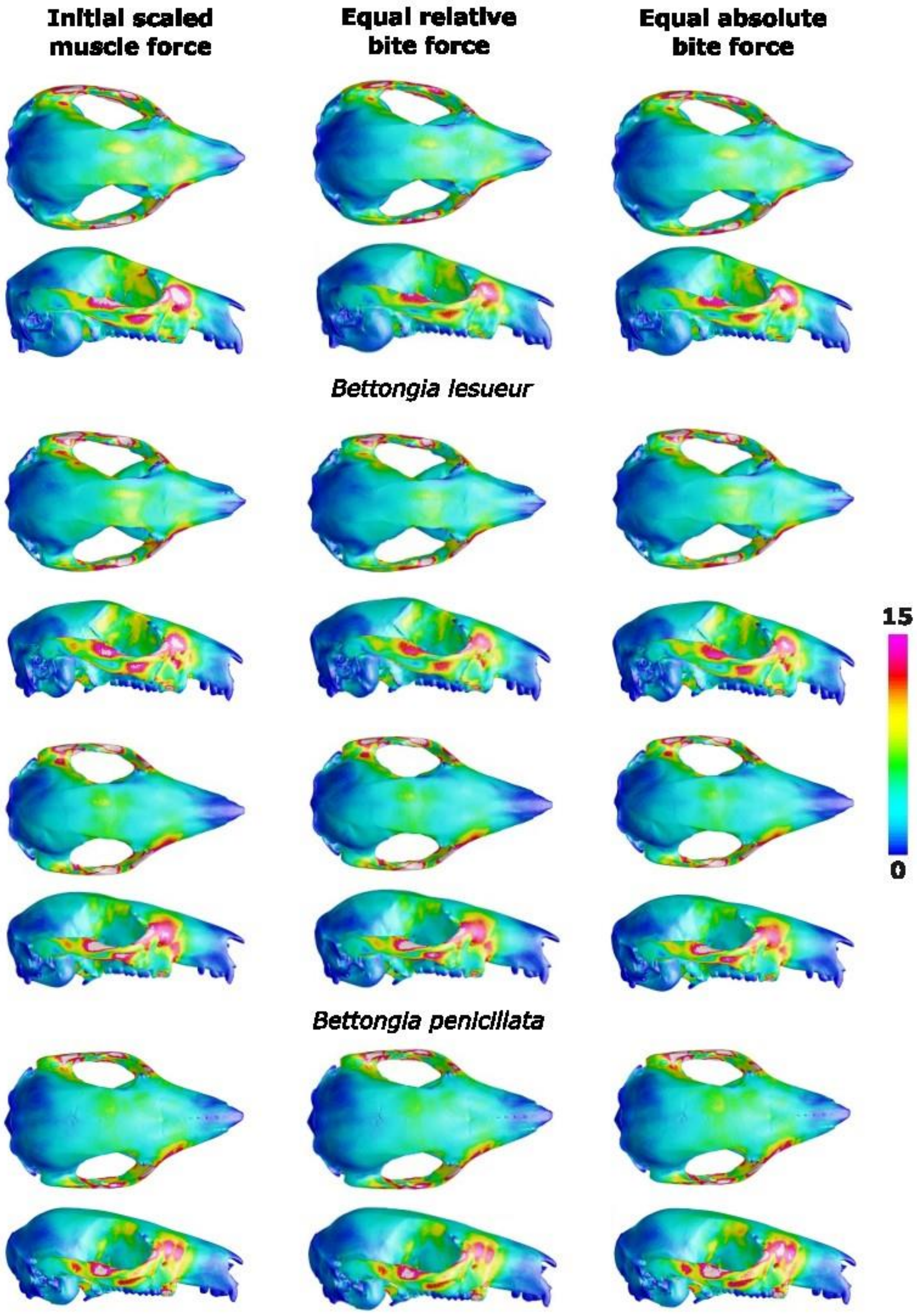

Initial scaled muscle force
Equal relative bite force
Equal absolute bite force
Bettongia lesueur
Bettongia penicillata
15
0

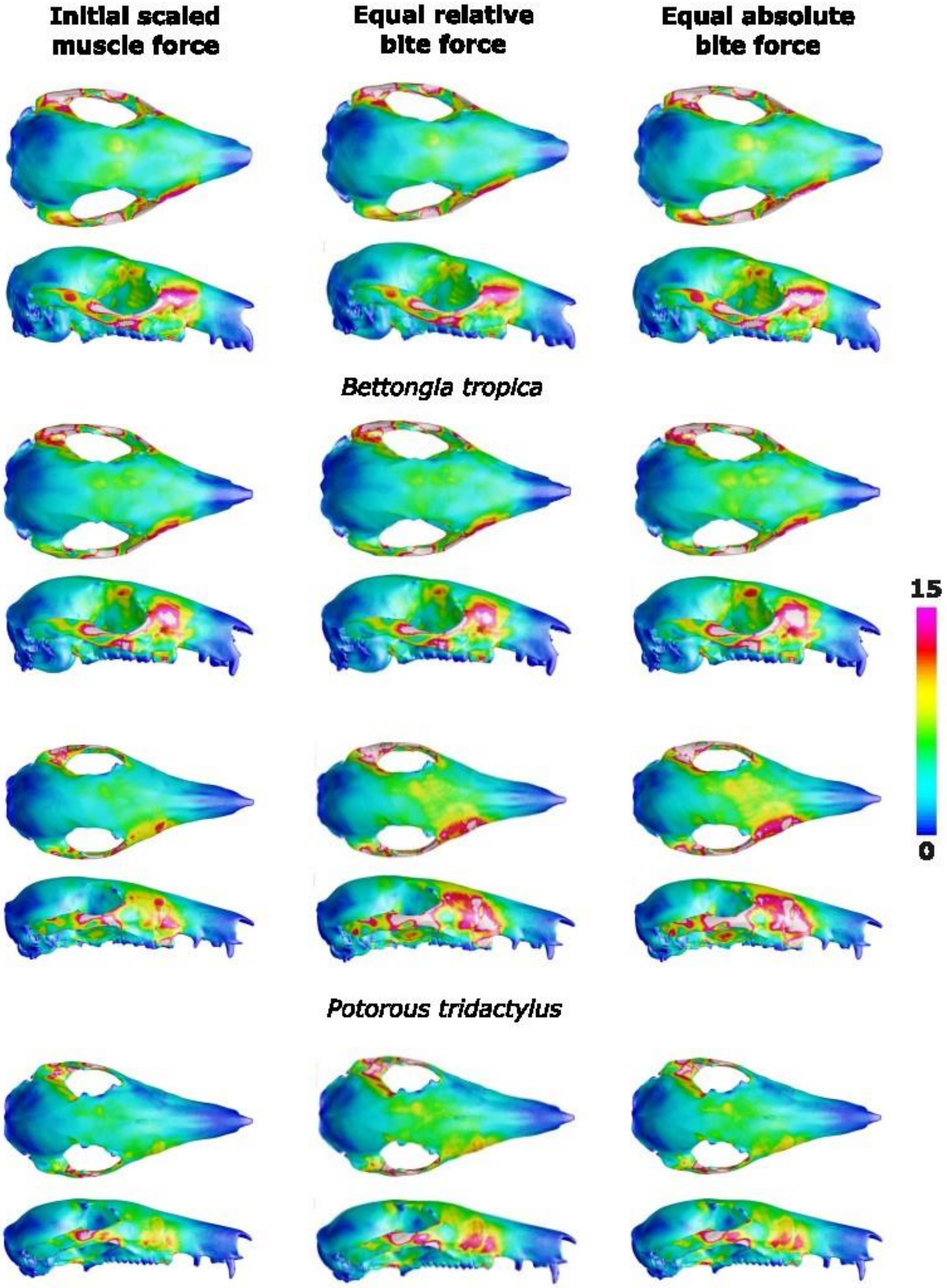
Initial scaled muscle force
Equal relative bite force
Equal absolute bite force
Bettongia tropica
15
0
Potorous tridactylus

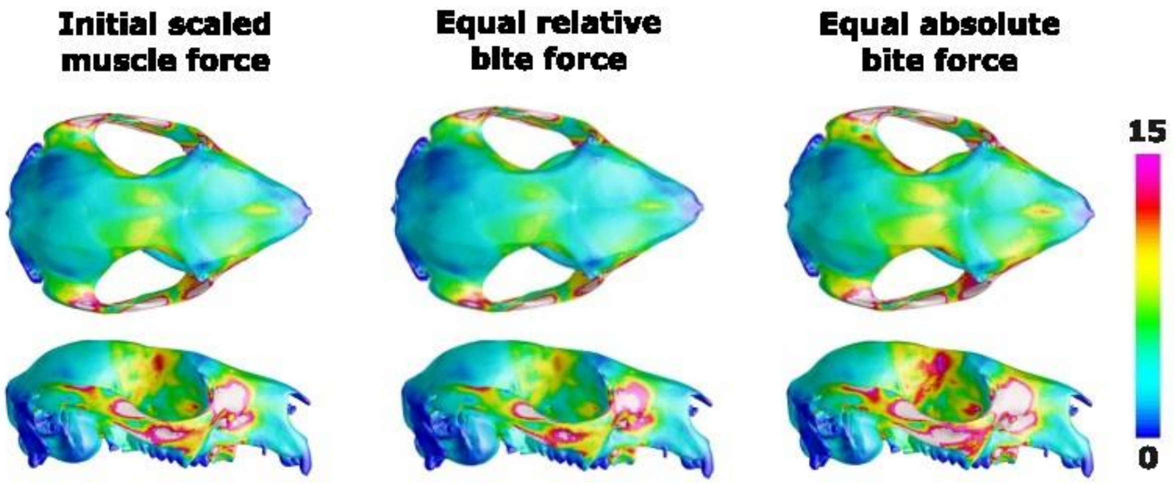


*Caloprymnus campestris*

Molar bite simulations

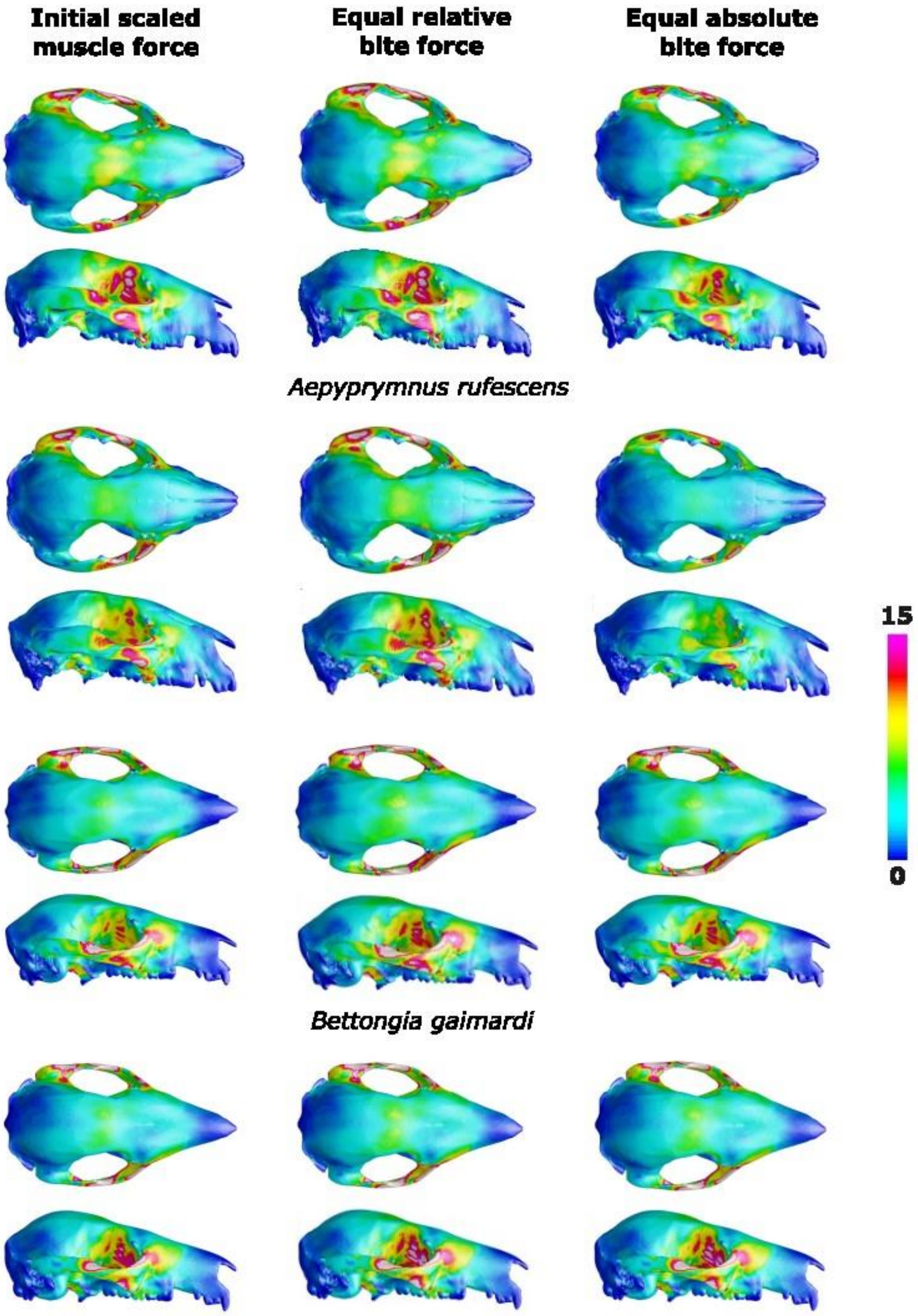

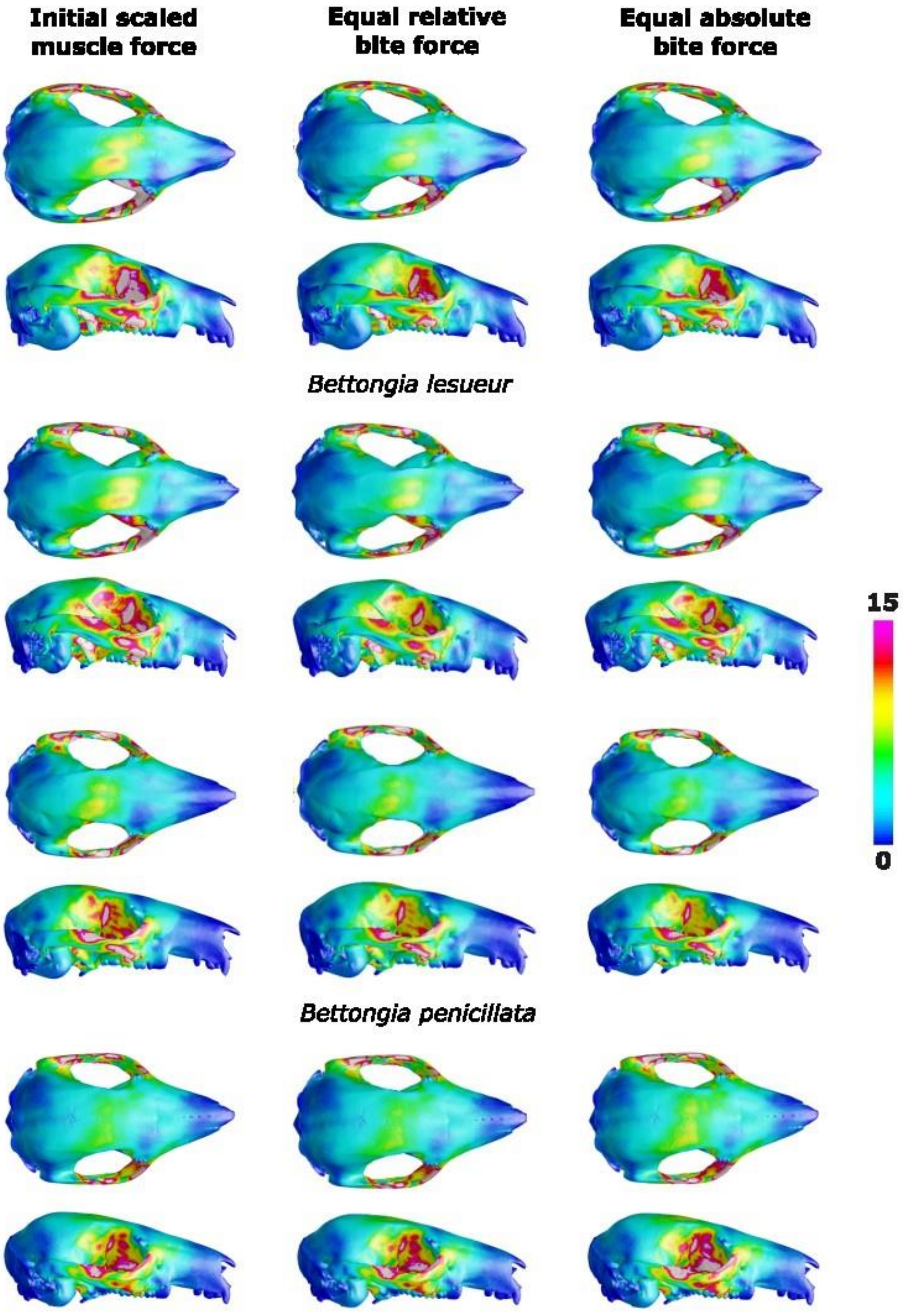
Initial scaled muscle force
Equal relative bite force
Equal absolute bite force
Bettongia lesueur
Bettongia penicillata
15
0

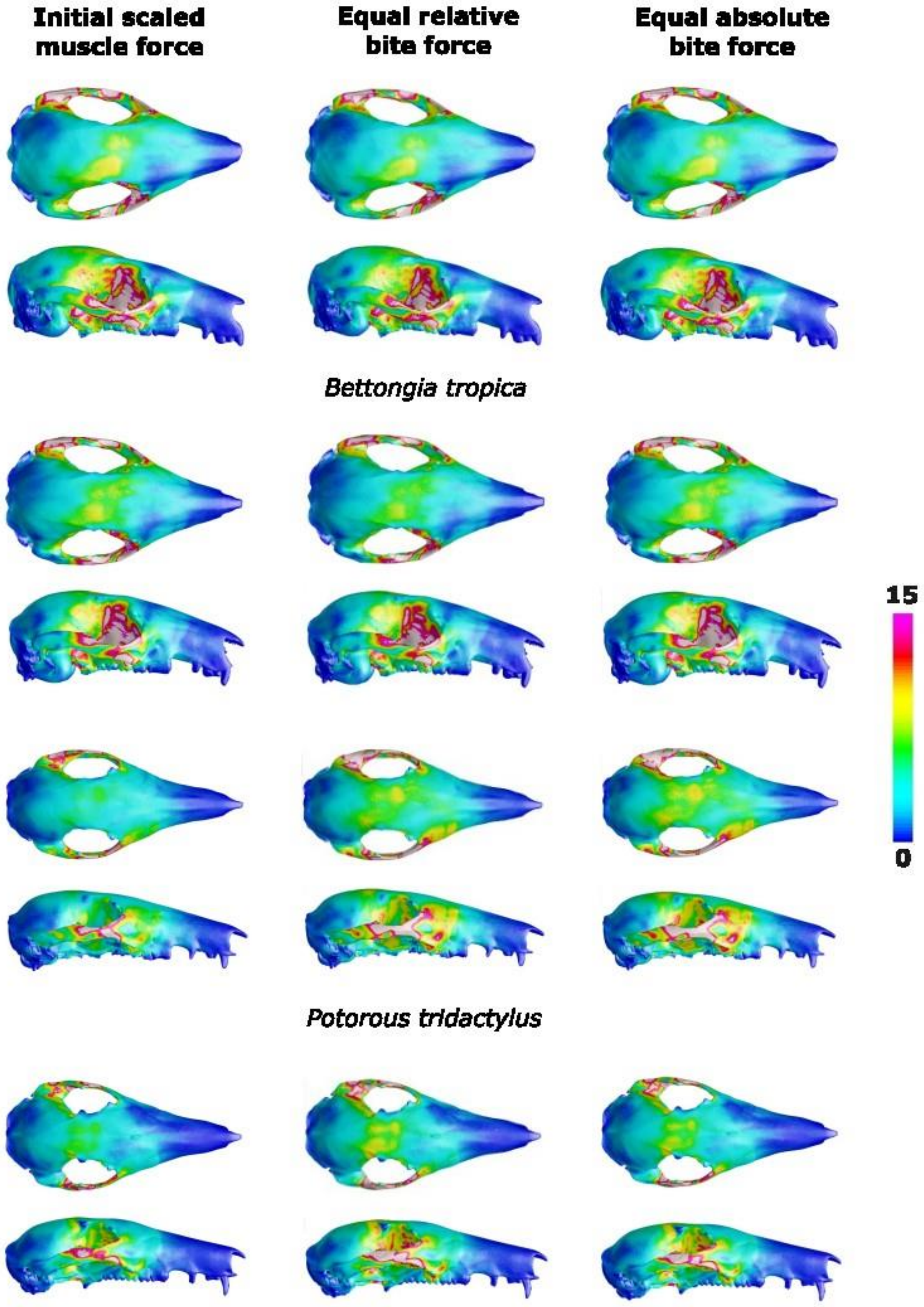
Initial scaled muscle force
Equal relative bite force
Equal absolute bite force
Bettongia tropica
15
0
Potorous tridactylus

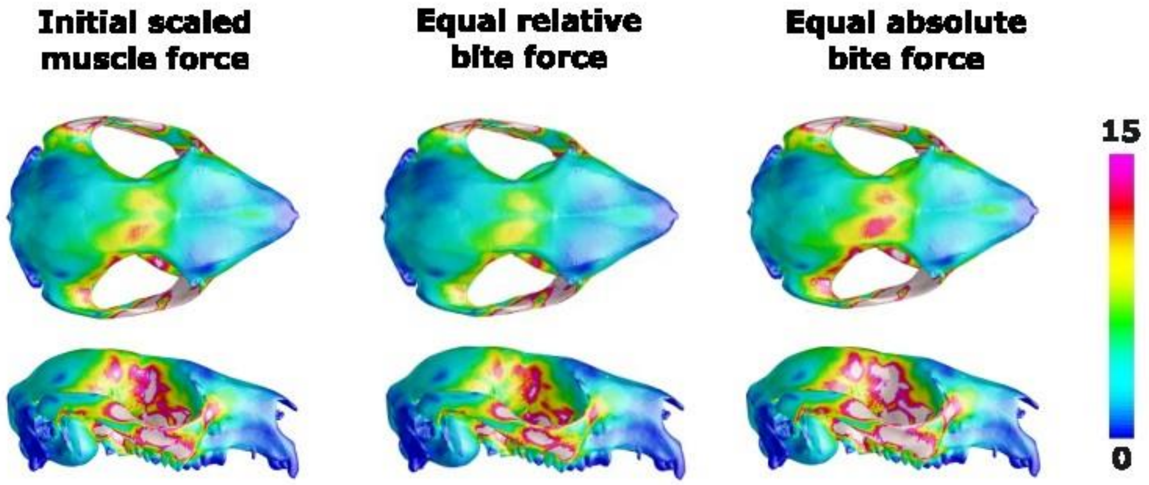


*Caloprymnus campestris*
